# The impacts of anesthetic regimens on the middle cerebral artery occlusion outcomes in male rats

**DOI:** 10.1101/2022.02.14.480371

**Authors:** Seyedeh Maryam Mousavi, Saeideh Karimi-Haghighi, Sara Chavoshinezhad, Sareh Pandamooz, Ivaldo Jesus Almeida Belém-Filho, Somaye Keshavarz, Mahnaz Bayat, Etrat Hooshmandi, Abbas Rahimi Jaberi, Mohammad Saied Salehi, Afshin Borhani-Haghighi

**Affiliations:** Clinical Neurology Research Center, Shiraz University of Medical Sciences, Shiraz, Iran; Cellular and Molecular Research Center, Research Institute for Health Development, Kurdistan University of Medical Sciences, Sanandaj, Iran; Stem cells Technology Research Center, Shiraz University of Medical Sciences, Shiraz, Iran; Department of Pharmacology, School of Medicine of Ribeirão Preto, University of São Paulo, Ribeirão Preto, São Paulo, Brazil; Department of Physiology, School of Medicine, Shiraz University of Medical Sciences, Shiraz, Iran

## Abstract

**Objectives:** The middle cerebral artery occlusion (MCAO) model was introduced more than three decades ago to simulate human stroke. Till now, it is the most common platform to investigate stroke-induced pathological changes as well as discover new drugs and treatments. Induction of general anesthesia is mandatory to induce this model, and different laboratories are using various anesthetic drugs, which might affect MCAO results. Therefore, the present study was designed to compare the impacts of several widely used anesthetic regimens on the MCAO outcomes.

**Materials and Methods:** Here, adult male rats were anesthetized by isoflurane inhalation, intraperitoneal injection of chloral hydrate, intraperitoneal injection of ketamine-xylazine, or subcutaneous administration of ketamine-xylazine, then subjected to 30 min MCAO. Mortality rate, body weight change, infarct size, as well as cognitive and neurological performance were evaluated up to three days after the surgery.

**Results:** Our findings revealed chloral hydrate caused the highest, while subcutaneous ketamine-xylazine led to the lowest mortality rate. Meanwhile, there were no significant differences in the body weight loss, infarct size, cognitive impairments, and neurological deficits among the experimental groups.

**Conclusions:** Based on the current results, we proposed that subcutaneous injection of ketamine-xylazine could be an effective anesthetic regimen in the rat model of MCAO with several advantages such as low mortality, cost-effectiveness, safety, ease of administration, and not requiring specialized equipment.

## Introduction

Stroke is among the major causes of death and inabilities around the world ^1, 2^. Conventional approaches such as thrombolysis and mechanical thrombectomy have transfigured the treatment of ischemic stroke as the primary type of stroke; however, the application of these therapies has been limited due to the risk of hemorrhage, narrow time windows, treatment failure, and availability ^3, 4^. Therefore, more preclinical investigations are required to discover next generations of drugs and treatments.

Animal models of stroke are fundamental tools for investigating stroke-induced pathological changes, as well as assessing the effectiveness and efficacy of potential therapeutic candidates. Up to now, different small and large animals such as mice, rats, rabbits, gerbils, cats, dogs, pigs, and monkeys have been used in stroke research ^5, 6^. Rat is one the ideal animals for simulating stroke due to their cerebrovascular similarity to humans, the capability to employ a variety of behavioral and neurological assessments after stroke, and moderate body size along with small brain size that facilitates the evaluation of physiological and pathological parameters ^7^.

Over the past decades, different approaches have been introduced to induce stroke in animal models, including middle cerebral artery occlusion (MCAO), photo-thrombosis, endothelin-1, embolic, fresh clot, lipid microparticle model, and platelet aggregation by using collagen fibril or adenosine diphosphate ^7^. Since MCA is the most frequently affected cerebral vessel in human ischemic cerebral vascular disease, the transient MCAO model is the most common platform for stroke research which is induced by occlusion of the MCA. However, various factors including strain, gender, weight, age, anatomical and vascular variation, filament characteristics, duration of ligation, collateral blood flow, follow-up period, and type of anesthesia can affect the stroke outcomes in experimental MCAO animals ^8, 9^.

In preclinical investigations, induction of MCAO requires general anesthesia, and a wide range of anesthetic drugs both injectable and inhalational, have been used for this purpose. Anesthesia can influence brain metabolism, inflammation, apoptosis, blood flow, cardiac output, and cerebral perfusion ^10, 11^, which interfere with experimental outcomes. Among the wide range of anesthetics, chloral hydrate (Table 2), isoflurane (Table 3), and ketamine/xylazine (Table 4) are the most commonly used drugs for the induction of anesthesia in stroke research.

**Table 2:**
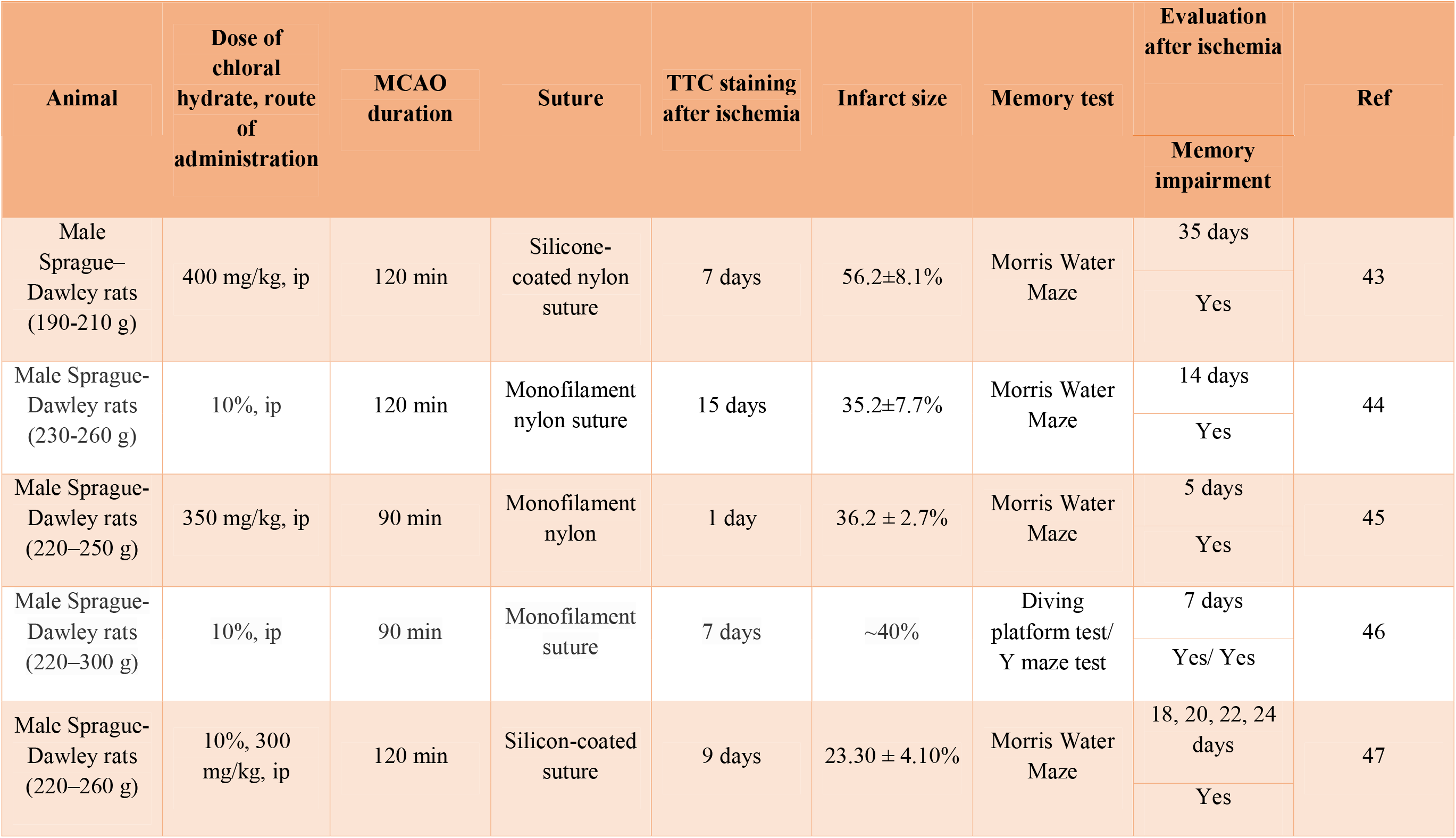
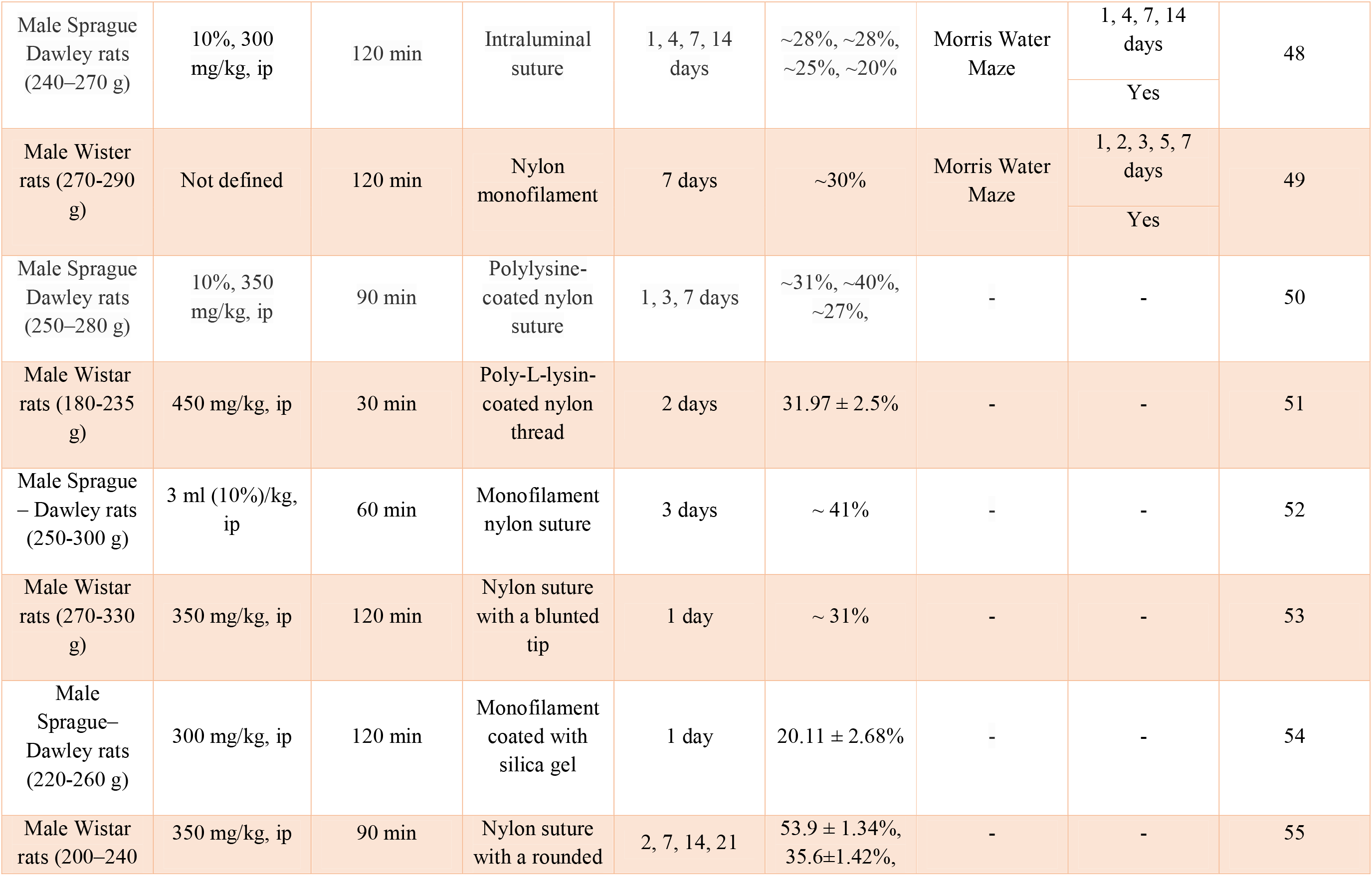
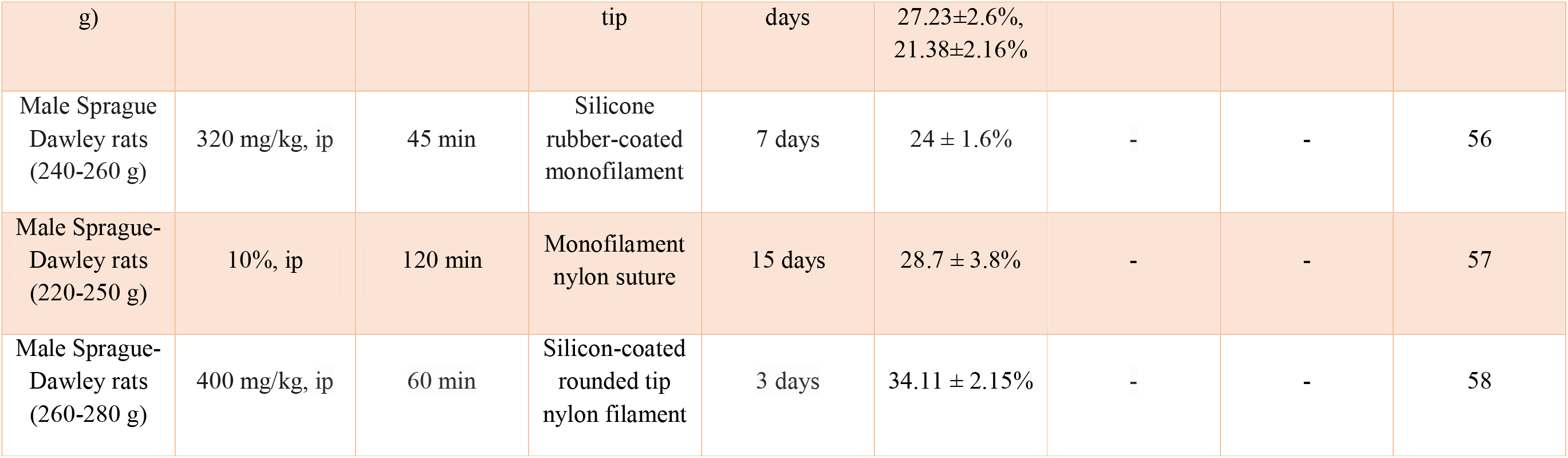
Infarct size and memory impairments in MCAO rats under chloral hydrate anesthesia.

| Animal | Dose of chloral hydrate, route of administration | MCAO duration | Suture | TTC staining after ischemia | Infarct size | Memory test | Evaluation after ischemia | Ref |
| --- | --- | --- | --- | --- | --- | --- | --- | --- |
|  |  |  |  |  |  |  | Memory impairment |  |
| Male Sprague–Dawley rats (190-210 g) | 400 mg/kg, ip | 120 min | Silicone-coated nylon suture | 7 days | 56.2±8.1% | Morris Water Maze | 35 days | 43 |
|  |  |  |  |  |  |  | Yes |  |
| Male Sprague-Dawley rats (230-260 g) | 10%, ip | 120 min | Monofilament nylon suture | 15 days | 35.2±7.7% | Morris Water Maze | 14 days | 44 |
|  |  |  |  |  |  |  | Yes |  |
| Male Sprague-Dawley rats (220–250 g) | 350 mg/kg, ip | 90 min | Monofilament nylon | 1 day | 36.2 ± 2.7% | Morris Water Maze | 5 days | 45 |
|  |  |  |  |  |  |  | Yes |  |
| Male Sprague-Dawley rats (220–300 g) | 10%, ip | 90 min | Monofilament suture | 7 days | ~40% | Diving platform test/ Y maze test | 7 days | 46 |
|  |  |  |  |  |  |  | Yes/ Yes |  |
| Male Sprague-Dawley rats (220–260 g) | 10%, 300 mg/kg, ip | 120 min | Silicon-coated suture | 9 days | 23.30 ± 4.10% | Morris Water Maze | 18, 20, 22, 24 days | 47 |
|  |  |  |  |  |  |  | Yes |  |
| Male Sprague Dawley rats (240–270 g) | 10%, 300 mg/kg, ip | 120 min | Intraluminal suture | 1, 4, 7, 14 days | ~28%, ~28%, ~25%, ~20% | Morris Water Maze | 1, 4, 7, 14 days | 48 |
|  |  |  |  |  |  |  | Yes |  |
| Male Wister rats (270-290 g) | Not defined | 120 min | Nylon monofilament | 7 days | ~30% | Morris Water Maze | 1, 2, 3, 5, 7 days | 49 |
|  |  |  |  |  |  |  | Yes |  |
| Male Sprague Dawley rats (250–280 g) | 10%, 350 mg/kg, ip | 90 min | Polylysine-coated nylon suture | 1, 3, 7 days | ~31%, ~40%, ~27%, | - | - | 50 |
| Male Wistar rats (180-235 g) | 450 mg/kg, ip | 30 min | Poly-L-lysine-coated nylon thread | 2 days | $31.97 \pm 2.5\%$ | - | - | 51 |
| Male Sprague – Dawley rats (250-300 g) | 3 ml (10%)/kg, ip | 60 min | Monofilament nylon suture | 3 days | ~ 41% | - | - | 52 |
| Male Wistar rats (270-330 g) | 350 mg/kg, ip | 120 min | Nylon suture with a blunted tip | 1 day | ~ 31% | - | - | 53 |
| Male Sprague–Dawley rats (220-260 g) | 300 mg/kg, ip | 120 min | Monofilament coated with silica gel | 1 day | $20.11 \pm 2.68\%$ | - | - | 54 |
| Male Wistar rats (200–240 | 350 mg/kg, ip | 90 min | Nylon suture with a rounded | 2, 7, 14, 21 | $53.9 \pm 1.34\%$, $35.6 \pm 1.42\%$, | - | - | 55 |
| g) |  |  | tip | days | 27.23±2.6%,<br>21.38±2.16% |  |  |  |
| Male Sprague<br>Dawley rats<br>(240-260 g) | 320 mg/kg, ip | 45 min | Silicone<br>rubber-coated<br>monofilament | 7 days | 24 ± 1.6% | - | - | 56 |
| Male Sprague-Dawley rats<br>(220-250 g) | 10%, ip | 120 min | Monofilament<br>nylon suture | 15 days | 28.7 ± 3.8% | - | - | 57 |
| Male Sprague-Dawley rats<br>(260-280 g) | 400 mg/kg, ip | 60 min | Silicon-coated<br>rounded tip<br>nylon filament | 3 days | 34.11 ± 2.15% | - | - | 58 |

**Table 3:** Infarct size and memory impairments in MCAO rats under isoflurane anesthesia.

| Animal | Dose of isoflurane | MCAO duration | Suture | TTC staining after ischemia | Infarct size | Memory test | Evaluation after ischemia | Ref |
| --- | --- | --- | --- | --- | --- | --- | --- | --- |
|  |  |  |  |  |  |  | Memory impairment |  |
| Male Sprague-Dawley rats (200-250 g) | 2% | 180 min | Nylon monofilament with a silicon rubber tip | 14 days | 41.8 ± 8.5% | Morris water maze | 14 days | 59 |
|  |  |  |  |  |  |  | Yes |  |
| Male Sprague-Dawley rats (250–70 g) | 2% in a mixture of 70% N2O and 30% O2 | 90 min | Monofilament nylon suture with rounded tip | 7 days | 19.48 ± 4.30% | Morris water maze | 14 days | 60 |
|  |  |  |  |  |  |  | Yes |  |
| Male Wistar rats (290–330 g) | 2% in a mixture of 50% N2O and 50% O2 | 90 or 120 min | Silicon rubber covered filament | 14 days | 90 min: ~13%<br>120 min: ~25% | Passive avoidance test/<br>Y-Maze test | 12 days/ 14 days<br>Yes/ No | 61 |
| Male Sprague–Dawley rats (270–320 g) | 1.5–2 %, in a mixture of N2O and O2 | 120 min | Poly-L-lysine coated monofilament | 1 day | ~35% | Morris Water Maze/ Novel Object Recognition | 28 days<br>Yes/ Yes | 62 |
| Male Sprague-Dawley rats (280-320 g) | 2% in a mixture of N2O and O2 | 90 min | Nylon thread tip | 5 days | ~350mm3 | Morris Water Maze | 28 days | 63 |
|  |  |  |  |  |  |  | No |  |
| Male Sprague-Dawley rats (250-300 g) | 1.5% in a mixture of 70% N<sub>2</sub>O and 30% O<sub>2</sub> | 45 min | Rounded tip surgical suture | 7 days | ~250mm<sup>3</sup> | - | - | 64 |
| Male Sprague-Dawley rats (270-300 g) | 1.5% in a mixture of 70% N<sub>2</sub>O and 30% O<sub>2</sub> | 90 min | Surgical suture | 15 days | 53.86 ± 8.60% | - | - | 65 |
| Male Sprague-Dawley rats (280±10 g) | 1% in a mixture of 70% N<sub>2</sub>O and 30% O<sub>2</sub> | 90 min | Rounded tip nylon suture | 1 day | 29.7 ± 5.2% | - | - | 66 |
| Male Sprague-Dawley rats (280-310 g) | 2% in a mixture of 70% N<sub>2</sub>O and 30% O<sub>2</sub> | 90 min | Silicone rubber-coated monofilament nylon suture | 1 day | 15.1±1.4% | - | - | 67 |
| Male Sprague-Dawley rats (280-320 g) | 1-3% in a mixture of 70% N<sub>2</sub>O and 30% O<sub>2</sub> | 90 min | Silica gel – coated nylon suture | 2 days | 42.52 ± 1.51% | - | - | 68 |
| Male Sprague-Dawley rats (180-220 g) | 2% in a mixture of 70% N<sub>2</sub>O and 30% O<sub>2</sub> | 90 min | Monofilament nylon suture with a rounded tip | 1 day | 40 ± 9% | - | - | 69 |
| Male Sprague- | 1% in a mixture of | 120 min | Poly-L-lysine-coated 4.0 | 3 days | 42.5% | - | - | 70 |
| Dawley Rats<br>(280–300 g) | 70% N2O and<br>30% O2 |  | nylon sutures |  |  |  |  |  |
| Male Sprague<br>–Dawley rats<br>(250–300 g) | 1.5% in a<br>mixture of<br>70% N2O and<br>30% O2 | 45 min | Rounded tip<br>surgical suture | 15-18 days | ~250 mm3 | - | - | 71 |

**Table 4:** Infarct size and memory impairments in MCAO rats under ketamine-xylazine anesthesia.

| Animal | Dose of ketamine-xylazine, Route of administration | MCAO duration | Suture | TTC staining after ischemia | Infarct size | Memory test | Evaluation after ischemia | Ref |
| --- | --- | --- | --- | --- | --- | --- | --- | --- |
|  |  |  |  |  |  |  | Memory impairment |  |
| Male Wistar rats (220–250 g) | ketamine (80 mg/kg) + xylazine (10 mg/kg) | 60 min | Poly-L-lysine-coated nylon monofilament | 1 day | ~180mm | Y-Maze | 1 day<br>Yes | 72 |
| Male Sprague–Dawley (260–280 g) | ketamine (80 mg/kg) + xylazine (10 mg/kg), ip | Permanent MCAO | Nylon suture with a silicon tip | 3, 4, 10 days | ~20% | Morris Water Maze | 30 days | 73 |
|  |  |  |  |  |  |  | Yes |  |
| Male Sprague-Dawley rats (250–300 g) | ketamine (80 mg/kg) + xylazine (10 mg/kg), ip | 120 min | Silicon coated nylon suture | 1 day | ~17% | Morris Water Maze | 25 days<br>Yes | 74 |
| Male Sprague–Dawley rats (240- 280 g) | ketamine (80 mg/kg) + xylazine (10 mg/kg), ip | 120 min | Silicon coated nylon suture | 1, 3 days | ~20% | Morris Water Maze | 28 days<br>Yes | 75 |
| Male Sprague-Dawley rats (250-300 g) | 60 mg/ml ketamine + 10 mg/ml xylazine (0.15 ml/100 g), ip | 90 min | Nylon monofilament surgical suture | 1 day | 29 ± 5% | - | - | 76 |
| Male Sprague-Dawley rats (250–300 g) | 100 mg/kg ketamine + 10 mg/kg xylazine, ip | 30 min | Monofilament nylon culture | 1 day | $576.3 \pm 5.5 \text{ mm}^3$ | - | - | 77 |
| Male Wistar rats (250–300 g) | 27-30 mg/kg ketamine + 3.6-4 mg/kg xylazine, ip | 90 min | Silicone-coated nylon suture | 14 days | $30.5 \pm 6.6\%$ | - | - | 78 |
| Male Sprague-Dawley Rats | 100 mg/kg ketamine + 10 mg/kg xylazine, ip | 90 min | poly-lysine coated monofilament nylon suture | 1 and 7 days | ~ 38% | - | - | 79 |
| Male Wistar rats (380–420 g) | 100 mg/kg ketamine + 10 mg/kg xylazine, ip | 60 min | Guide wire with a blunted tip | 1 day | ~ 400 mm<sup>3</sup> | - | - | 38 |
| Female Sprague-Dawley rats (250–300 g) | 75 mg/kg ketamine + 10 mg/kg xylazine, ip | Permanent MCAO | Rounded tip surgical suture | 7 days | $253.91 \pm 8.51 \text{ mm}^3$ | - | - | 80 |
| Female Sprague-Dawley rats (250–300 g) | 50 mg/kg ketamine + 10 mg/kg xylazine, ip | Permanent MCAO | Rounded tip surgical suture | 15 days | ~ 200 mm<sup>3</sup> | - | - | 81 |
| Male Sprague– | 80 mg/kg ketamine + 40 | 120 min | Poly-l-lysine-coated | 1 day | ~ 25% | - | - | 82 |
| Dawley rats<br>(380–420 g) | mg/kg<br>xylazine, ip |  | monofilament<br>nylon suture<br>with the tip<br>rounded |  |  |  |  |  |
| Male<br>Sprague–<br>Dawley rats<br>(200–220 g) | 100 mg/kg<br>ketamine + 15<br>mg/kg<br>xylazine, ip | 120 min | Poly-l-lysine-coated<br>monofilament<br>nylon suture<br>with the tip<br>rounded | 1 day | ~55–75%<br>contralateral<br>hemisphere | - | - | 83 |

Chloral hydrate is one of the oldest sedatives and hypnotic medications with little or no analgesic effects. Also, it has anticonvulsant and muscle relaxant properties ^12^. Although the safety margin of chloral hydrate is narrow, it is traditionally used as an anesthetic drug in stroke studies ^13^. Isoflurane, a potent volatile anesthetic, is approved by the Federal Drug Administration for the induction and maintenance of anesthesia. Also, isoflurane reduces pains sensitivity and induces muscle relaxation, amnesia, as well as sedation, and currently, it has been widely used in stroke studies ^13^. Ketamine is an N-methyl-D-aspartate (NMDA) receptor antagonist, and xylazine is an α2-adrenergic agonist ^14^. For decades, a combination of ketamine-xylazine with a wide margin of safety has been used for general anesthesia in the experimental stroke models.

Considering the potential of anesthetics to affect stroke outcomes, we aimed to compare the effects of ketamine/xylazine, chloral hydrate, and isoflurane anesthetic drugs on the survival rate, body weight, immobilization time, induction time, recovery time, infarct size, cognitive ability, and neurological function of rats following MCAO induction.

## Material and methods

### Animals and ethics statement

Sixty adult male Sprague Dawley rats (weighting 240-260 g) were obtained from the Center of Comparative and Experimental Medicine, Shiraz University of Medical Sciences. All rats were maintained in a controlled condition with free access to standard food and water. The present study was approved by the Animal Care Committee of Shiraz University of Medical Sciences, Shiraz, Iran (IR.SUMS.REC.1400.452) and performed in compliance the National Research Council’s Guide for the Care and Use of Laboratory Animals. All attempts were conducted to minimize the number of animals and decrease animal suffering.

### Experimental design

All animals were randomly divided into four groups (15 rats in each group) with different anesthetic protocols as follows: 1- Anesthetized with an intraperitoneal injection of chloral hydrate (#C8383, Sigma-Aldrich, 320 mg/kg; CH group); 2- Anesthetized with isoflurane inhalation (Terrell, Piramal, 2%; Iso group) and 3- Anesthetized with an intraperitoneal injection of ketamine and xylazine (Alfasan, 100 and 5 mg/kg, respectively; Ket-Xyl/ IP group) (Figs. 1A and B). Moreover, previously it has been reported that subcutaneous administration of ketamine-xylazine in mice is as effective as the intraperitoneal route ^15^. Hence, in the present study, the fourth group was defined to anesthetize rats with subcutaneous injection of ketamine and xylazine (Alfasan, 100 and 5 mg/kg, respectively; Ket-Xyl/ SC group) (Fig. 1C). In each experimental group, animals were randomly subdivided into SHAM (n=5) and MCAO (n=10) rats.

**Figure1:**
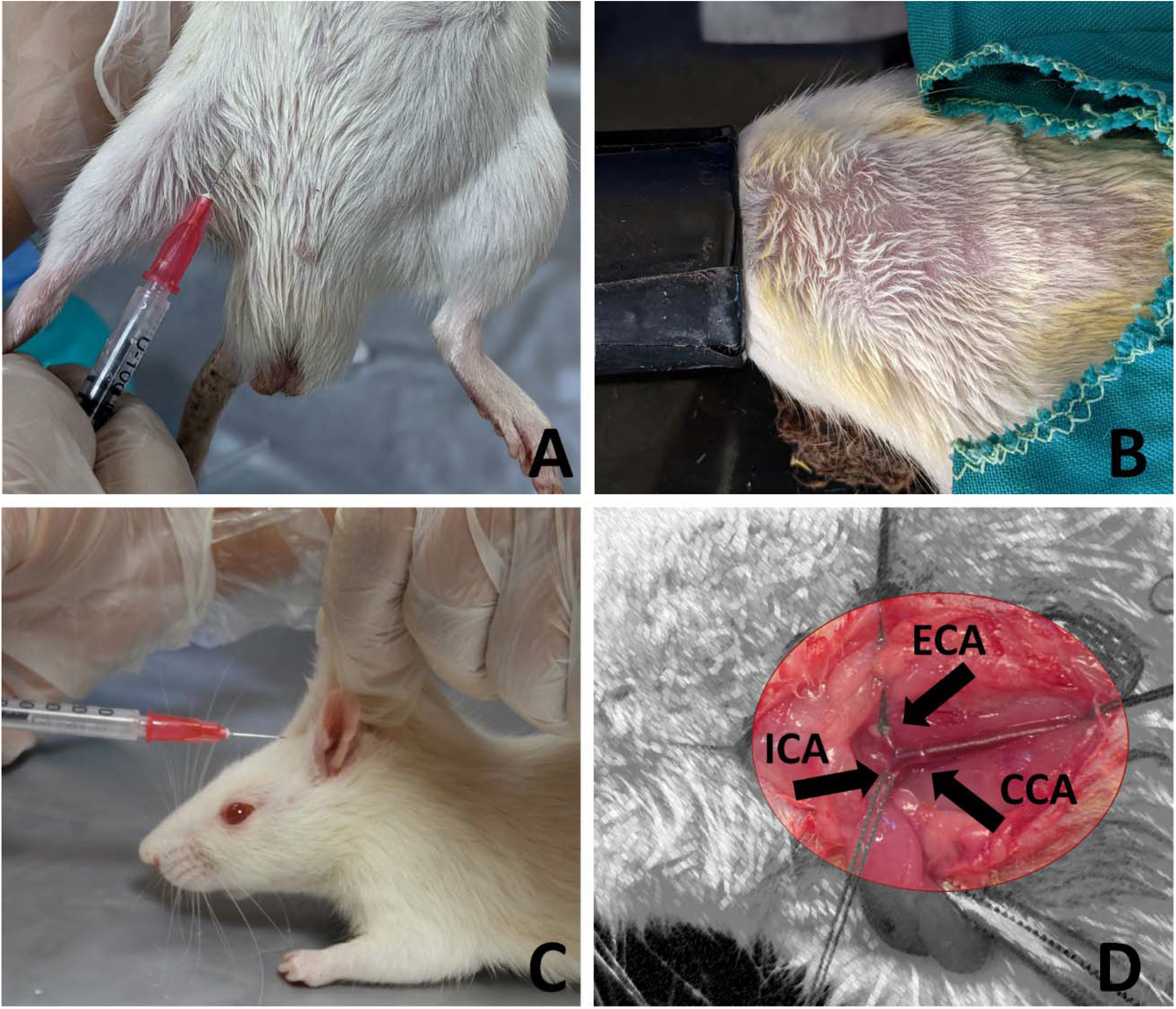
Different routes of anesthetics administration. A: Intraperitoneal, B: Inhalation, C: Subcutaneous. D: Induction of MCAO model based on the Koizumi’s method. CCA, common carotid artery; ECA, external carotid artery; ICA, internal carotid artery.

### Transient MCAO procedure

To induce ischemic stroke, rats underwent transient unilateral MCAO. Briefly, an incision was made in the middle of the neck, then right common carotid, external carotid, and pterygopalatine arteries were ligated. A small incision was made in the right common carotid artery, then a silicone-coated monofilament (#403556, Doccol Corporation) was inserted. The monofilament was cranially advanced to the internal carotid artery until a mild resistance was felt (Fig. 1D). After 30 minutes ^16, 17^, the filament was carefully removed to permit blood-flow reperfusion. Laser Doppler (ML191, AD Instrument, Australia) was applied to check microvascular blood flow decrease during the surgery. The SHAM rats underwent a similar surgical procedure without monofilament insertion.

In all experimental groups, time to onset of immobilization (from anesthetic administration to loss of righting reflex), time to onset of surgery (from anesthetic administration to loss of pedal reflex), and time from surgery to recovery (from loss of pedal reflex to gain it) were recorded.

### Evaluation of survival rate and body weight

Mortality was recorded daily, and obtained findings were presented as survival rates in each experimental group. Furthermore, body weight was recorded daily as a general marker for well-being.

### Evaluation of neurological deficits

All animals were evaluated for neurological deficits on day 0 before surgery as well as 1, and 3 days after the surgical procedure. The neurological function was graded on a scale of 0-4, as it is shown in Box 1.

### Evaluation of spatial short-term memory

The Y-maze test was carried out three days after the surgery to evaluate the short-term spatial working memory. The Y-maze apparatus was made of black Plexiglas with three arms (50 × 10 × 30 cm, named A, B, C), which oriented 120° angle to each other. Each rat was initially placed at the end of one arm and allowed to move freely through the maze, and the sequence of entered arms (with four paws) was recorded for 8 minutes. The total number of arm entries was considered as an index of locomotion. The rats which performed < 8 arm entries during the test were excluded. Moreover, spontaneous alternation performance (SAP) and alternate arm return (AAR) were calculated as follows:

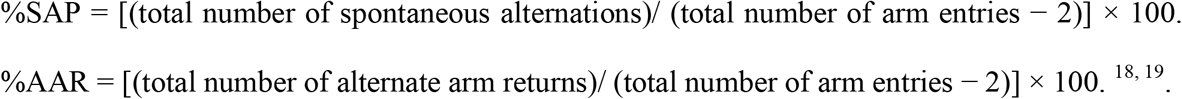

### Evaluation of infarct size

Three days following the surgery, all animals were killed to assess the infarct size using 2,3,5-triphenyltetrazolium chloride (TTC) staining. Under deep anesthesia with CO2, rats’ brains were removed and coronally cut (2mm) using a brain matrix. The brain sections were incubated in 0.5% TTC (Sigma) for 30 minutes at 37°C, and infarct size was calculated using ImageJ software.

### Statistical analysis

Statistical analysis and data graphing were performed using GraphPad Prism (Version 9.2, GraphPad Software, Inc). Neurological deficit scores were statistically analyzed using the nonparametric Kruskal-Wallis test and presented as median with interquartile interval. The %infarct size, and Y-maze data were subjected to the Shapiro-Wilk normality test, and comparisons among groups were made by one□way analysis of variance (ANOVA) followed by post hoc Tukey test. Two-way ANOVA repeated measure followed by post hoc Tukey test was used to analyze body weight and presented as mean ± SEM. The *P*< 0.05 was considered statistically significant.

## Results

### Survival rate

Survival of experimental rats was evaluated daily (Fig. 2). According to obtained data, no mortality occurred in SHAM groups throughout the experiment (Fig. 2a). On the other hand, cerebral ischemia resulted in different mortality rates in MCAO groups (Fig. 2A). Induction of ischemia in CH anesthetized rats led to the highest mortality (40%). The mortality rate in Ket-Xyl/ SC group was 20%, while this rate in isoflurane and Ket-Xyl / IP groups was 30%.

**Figure 2:**
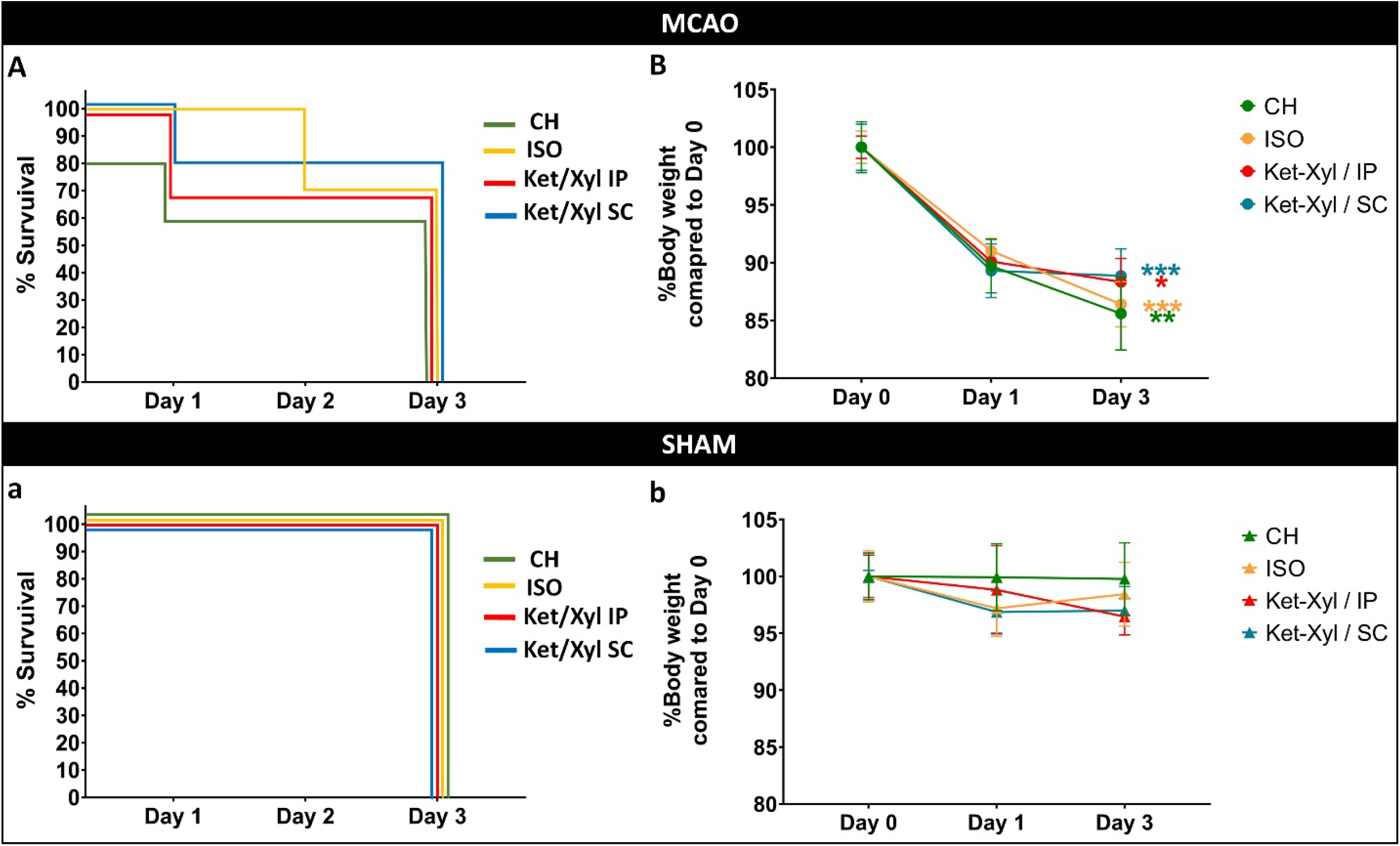
The effects of different anesthetic regimens on the mortality percentage and body weight changes in the MCAO (upper panel) and SHAM (lower panel) groups. *P < 0.05, **P < 0.01, and ***P < 0.001 significant differences between days 1 and 3 post-ischemia. Data are expressed as mean ± SEM.

### Body weight change

Body weight was assessed before, as well as 1 and 3 days post-surgery. A considerable weight loss was observed in all ischemic groups at day 3 compared to day 0 (Fig. 2B); however, the body weight changes in SHAM groups were not statistically significant at various time points (Fig. 2b).

### Functional deficits

In the current study, neurological function was evaluated before, as well as 1 and 3 days after the surgery. Before surgery, no deficit was detected in any groups. At 1 and 3 days post-MCAO, ischemic rats exhibited remarkable functional deficits, and no significant differences were detected among the four different anesthetic protocols (Figs. 3A & B). No deficit was observed in the SHAM groups at any time points (Figs. 3a & b).

**Figure 3:**
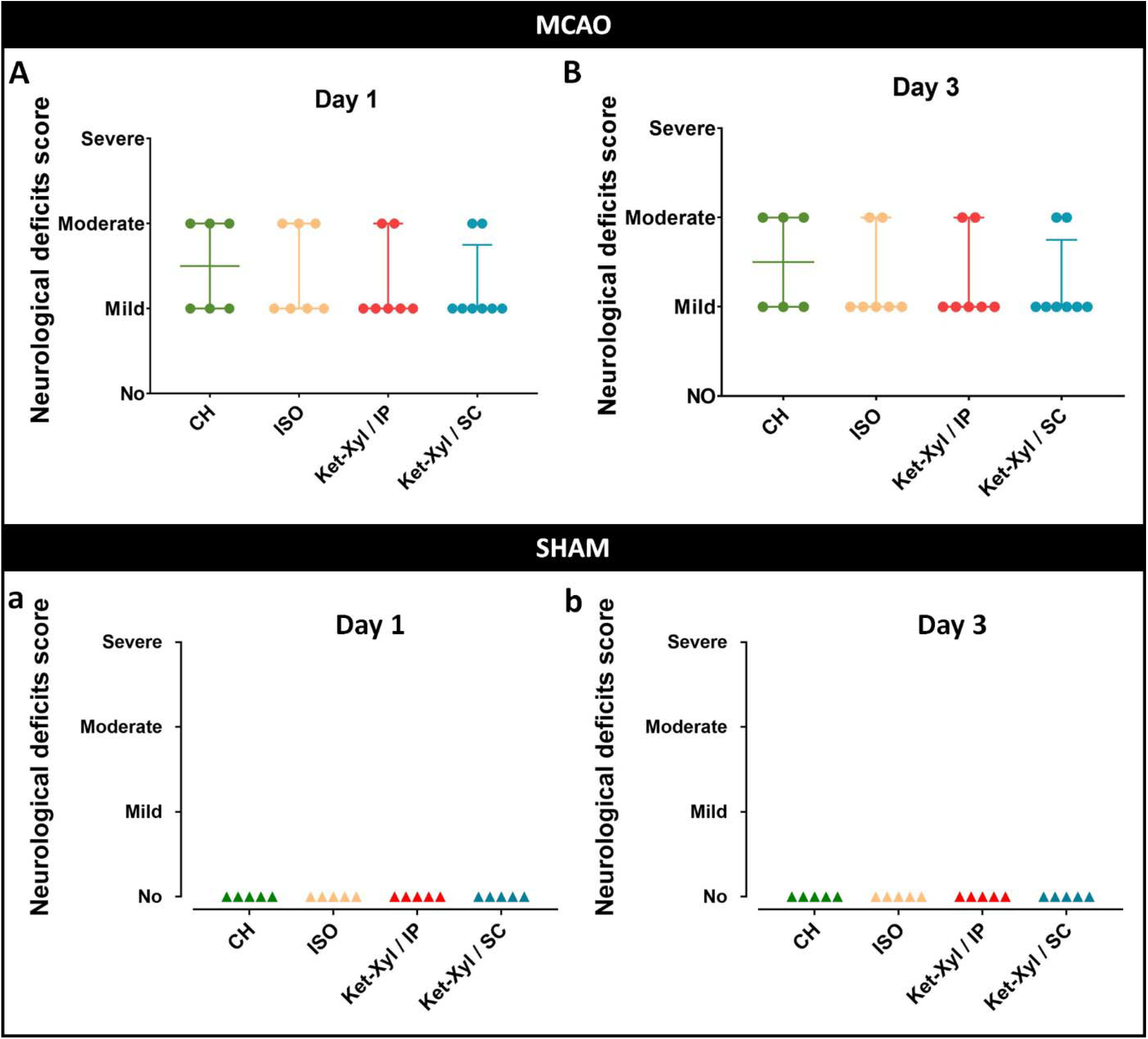
The effects of different anesthetic regimens on the neurological deficits at 1 and 3 days following surgery in MCAO (upper panel) and SHAM (lower panel) groups. Data are expressed as median ± interquartile interval. There was no significant difference between experimental groups.

### Time to onset of immobilization

The time to onset of immobilization (from anesthetic administration to loss of righting reflex) was ranged from 43±10s in isoflurane inhaled anesthesia to 76±7s in CH anesthetized rats. In Ket-Xyl/ IP and Ket-Xyl/ SC groups, this time was 53±15s and 62±17s, respectively. There was no significant difference in the time to onset of immobilization between all anesthetic regimens (Table 1).

**Table 1:** The impacts of different anesthetic regimens on the several surgical parameters. Different letters indicate significant difference (P<0.05).

| Experimental groups | Time to onset of immobilization | Time to onset of surgery | Time from surgery to recovery |
| --- | --- | --- | --- |
|  | (sec) | (min) | (h) |
| **CH** | $76 \pm 7$ | $6.3 \pm 0.5^b$ | $1.25 \pm 0.2$ |
| **ISO** | $43 \pm 10$ | $1.8 \pm 0.4^a$ | - |
| **Ket-Xyl / IP** | $53 \pm 15$ | $8.7 \pm 0.7^c$ | $1.65 \pm 0.2$ |
| **Ket-Xyl / SC** | $62 \pm 17$ | $8.5 \pm 0.4^c$ | $1.75 \pm 0.1$ |

### Time to onset of surgery

Obtained data revealed a significant difference regarding the average time to onset of surgery (from anesthetic administration to loss of pedal reflex). This time was ranged from 1.8±0.4 min for isoflurane inhaled anesthesia to 8.7±0.7 min in rats anesthetized with an intraperitoneal injection of Ket-Xyl. This period was 6.3±0.5 and 8.5±0.4 min in CH and Ket-Xyl/ SC groups, respectively (Table 1).

### Time from surgery to recovery

This time course was measured from loss of pedal reflex to gain it. This period was ranged from 1.25±0.2 h in the CH-anesthetized group to 1.75±0.1 h in the Ket-Xyl/ SC group. This duration was 1.65±0.2 h in rats anesthetized with an intraperitoneal injection of Ket-Xyl. There was no statistical difference among the three anesthetic-injected groups. Experimental rats that were anesthetized with isoflurane, recovered immediately after breathing circuit removal (Table 1).

### Infarct size

The infarct size was assessed by TTC staining three days after the surgery. Our data demonstrated dramatic MCAO-induced damages in the ipsilateral hemisphere of all ischemic groups (Fig. 4A). There was no significant difference in infarct size between all anesthetic regimens (Fig. 4B). No brain insult was detected in the SHAM groups (Figs. 4a & b).

**Figure 4:**
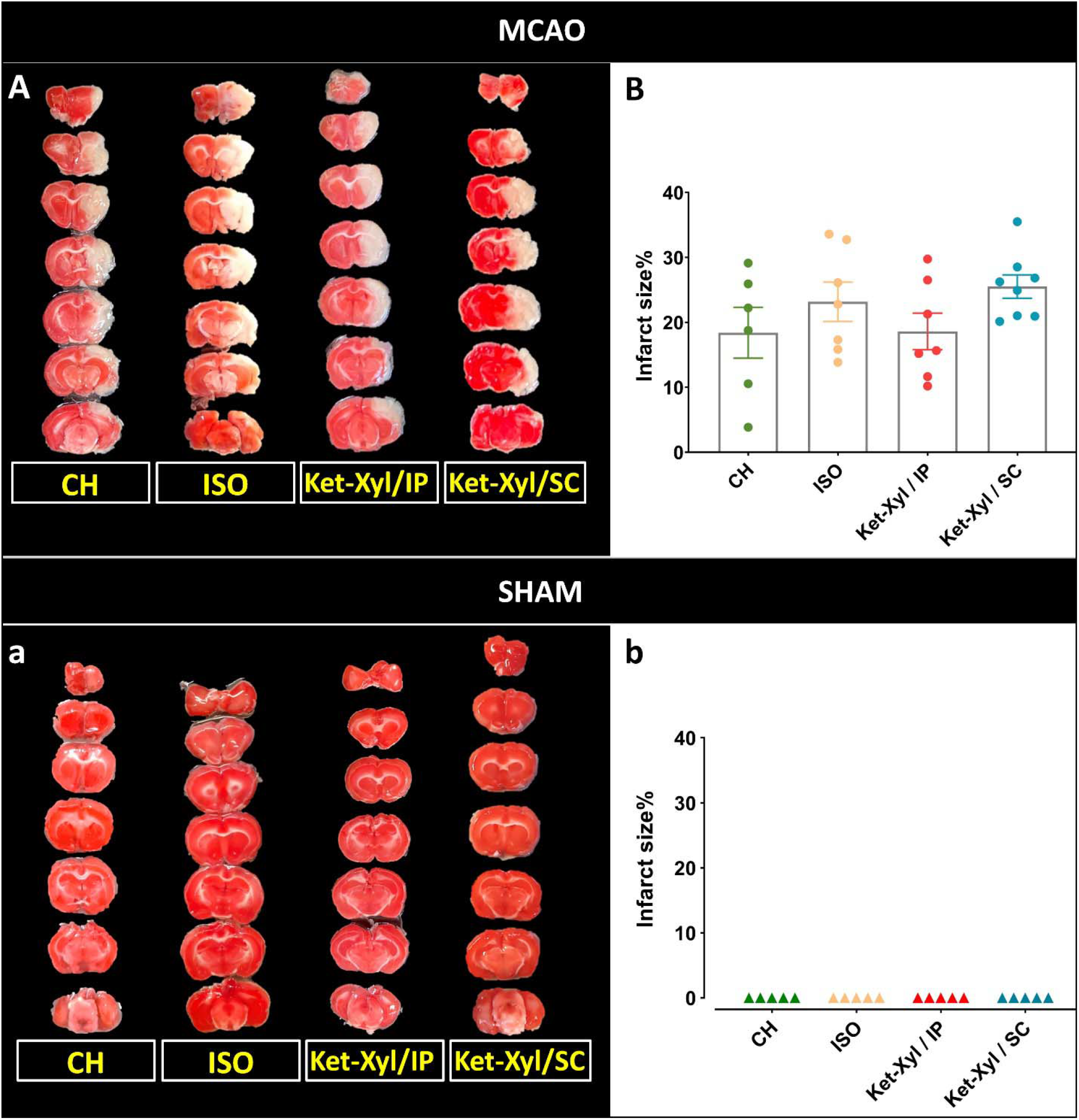
The effects of different anesthetic regimens on the infarct size. Coronal depiction of brain sections 3 days post-surgery in all experimental groups stained with TTC (A and a). Quantification of %infarct size in MCAO (B) and SHAM (b) groups. Data are expressed as mean ± SEM. There was no significant difference between experimental groups.

**Figure 5:**
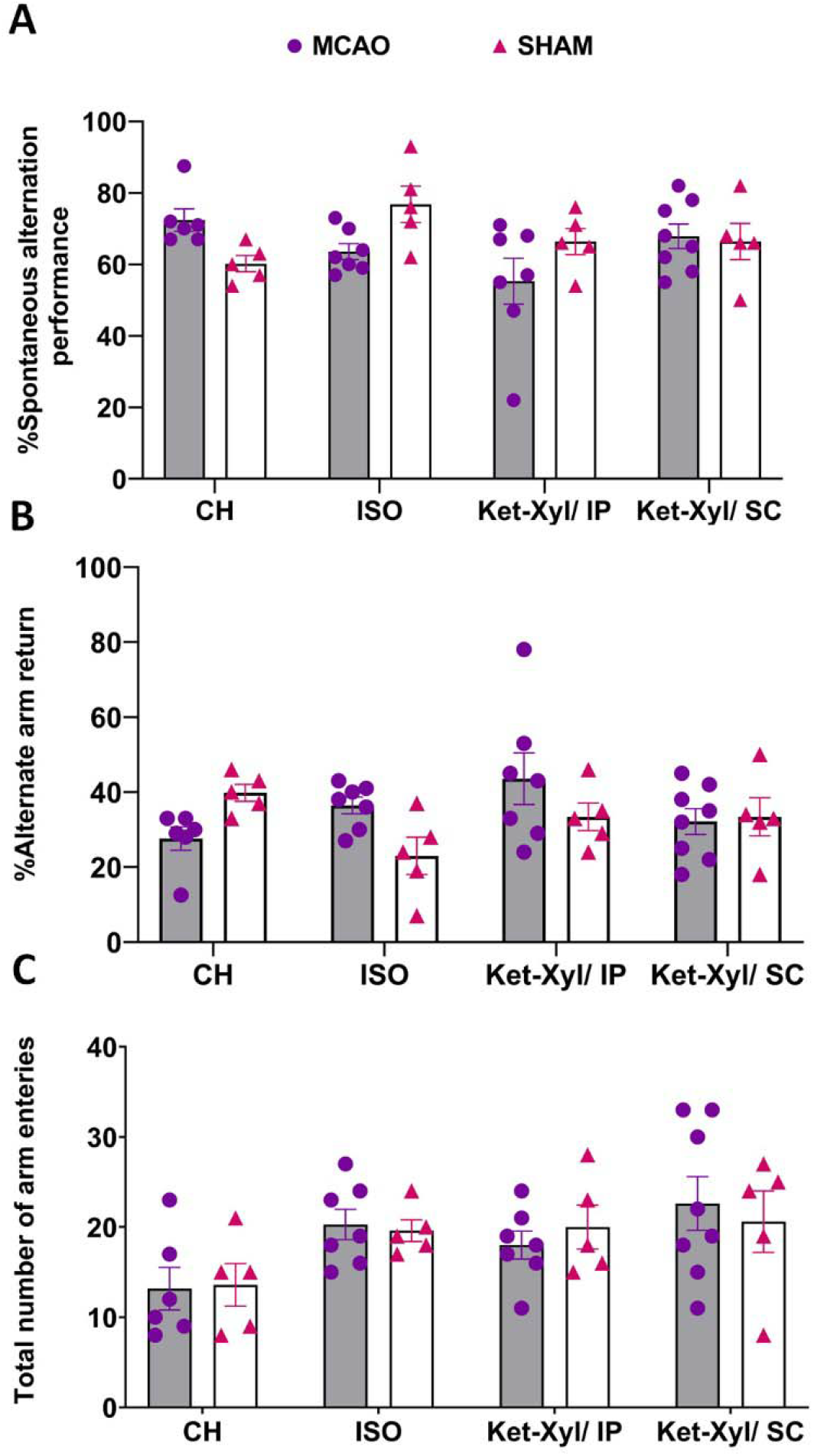
The effects of different anesthetic regimens on the spatial working memory as assessed by Y-maze test. A: Percentage of spontaneous alternation behavior (%SAP); B: Percentage of alternate arm return (%AAR); C: Total number of arm entries as an indicator of locomotor activity. Data are expressed as mean ± SEM. There was no significant difference between experimental groups.

### Spatial short-term memory

Our data indicated no significant difference among all experimental groups, including MCAO subdivided groups, and SHAM subdivided groups, neither in AAR%, nor SAP%. Although a significant difference was detected between the locomotor activity of the CH and Ket-Xyl/ SC ischemic rats, no statistical difference was observed between other groups.

## Discussion

Animal models of cerebral ischemia are valuable assets in the understanding stroke pathophysiology as well as the identification of novel therapeutic strategies ^20^. The MCAO model is one of the most commonly employed platforms that was developed more than three decades ago to simulate human ischemic stroke ^21^. Transient MCAO using intravascular filament in experimental animals, possesses several advantages such as reproducibility, not requiring craniectomy, and simply allowing reperfusion by the withdrawal of the occluding monofilament; however, induction of general anesthesia is mandatory to induce this model ^22^. It has been reported that anesthetics have potential impacts on the stroke outcome in a drug- and dose-dependent manner ^23^. Anesthetic agents could alter cerebral blood flow, neurovascular coupling, autoregulation, ischemic depolarizations, excitotoxicity, inflammation, and neural networks during surgery which confound the interpretation of stroke outcome ^10, 23^. Hence, the anesthetic regimen in experimental stroke models needs to be carefully selected and optimized. Safety, availability, cost-effectiveness, ease of administration, low mortality, minimal impact on physiological and functional parameters, and postoperative analgesic effect are the main factors that should be considered in stroke research ^13^.

Isoflurane is a potent inhalation anesthetic with minimal metabolism and organ toxicity. It reduces pain sensitivity and relaxes muscles by binding to gamma-aminobutyric acid (GABA), glutamate and glycine receptors. Isoflurane enhances intrasynaptic GABA-A mediated currents and activates glycine receptor in the central nervous system, which leads to decreased motor function ^24^. Moreover, it inhibits NMDA glutamate receptor and potassium channels in the spinal cord that participate in skeletal muscle relaxation ^25^. Isoflurane is currently used in stroke experimental studies due to its low mortality rate, as well as rapid induction and recovery kinetics. However, requiring specialized equipment, high cost, rapid consumption, and damaging environmental effects, has been limited its use compared to injectable anesthetics.

Among the various injectable anesthetics, chloral hydrate and ketamine-xylazine are also commonly used in MCAO studies. Chloral hydrate is a sedative-hypnotic agent; however, its precise mechanism of action is unknown. It has been previously shown that chloral hydrate is metabolized by the liver and erythrocytes to form active metabolite trichloroethanol. Trichloroethanol is an agonist for the GABA-A receptors, and is responsible for opening a chloride ion channel and depression of the central nervous system ^26^. Chloral hydrate can also enhance glycine receptor activity and facilitate 5-hydroxytryptamine-3 receptor-mediated currents in ganglionic neurons ^27^. Moreover, chloral hydrate inhibits α-amino-3-hydroxy-5-methyl-4-isoxazole-propionic acid (AMPA) glutamate receptors-induced calcium influx in neurons ^28^. Despite the traditional application of chloral hydrate in the MCAO animal model because of its minimal effects on stroke outcomes, its use is associated with a variety of disadvantages. Chloral hydrate has no analgesic effect and induces respiratory depression and cardiac dysrhythmias at surgical doses, which lead to an increase in mortality rate in animals. Moreover, chloral hydrate is carcinogenic and could cause ileus, peritonitis, and gastritis ^29, 30^.

Ketamine, a water-soluble phencyclidine derivative, is a noncompetitive NMDA receptor antagonist that blocks the phencyclidine sites on NMDA receptors in the cerebral cortex, hippocampus, and spinal cord, resulting in neuronal depolarization ^31^. Moreover, ketamine can modulate the activity of opioid receptors, monoamine, cholinergic, purinergic, and adrenoreceptor systems ^32^. Ketamine has a wide margin of safety and induces amnesia, analgesia, and immobility. In addition to its anesthetic effects, ketamine has been shown to induce catalepsy, somatic analgesia, bronchodilation, and sympathetic nervous system stimulation ^32^. In peripheral tissues, ketamine is mainly metabolized into nor-ketamine through hydroxylation and N-demethylation. Nor-ketamine is the primary active metabolite of ketamine ^33^. For anesthesia, ketamine is combined with other agents like xylazine (α2-adrenergic agonist) to achieve mild to moderate analgesia, and muscle relaxation ^34^. Ketamine-xylazine anesthesia preserves blood pressure and spontaneous respiratory function better than chloral hydrate ^35^. Ketamine and xylazine are frequently used for anesthesia in animal models of ischemia.

The mixture of ketamine-xylazine is usually injected into the peritoneal cavity. Despite rapid absorption of drugs, the intraperitoneal administration has some limitations, including pain and irritation at the site of injection, formation of fibrous tissue within the abdominal cavity, risk of hemorrhage, along requiring safe restraint of the animal. Furthermore, intraperitoneal administration might increase the release of inflammatory cytokines from mesenteric arteries, which can interfere with the results ^15, 35^. Subcutaneous administration provides a safe, less painful, less invasive, and easy route of drug infusion. In this route, the drug is slowly absorbed by small capillaries in the skin, which in turn provides a long-term effect ^15, 36^. Subcutaneous ketamine administration has been used to control human pain ^37^. In addition, it has been reported that subcutaneous injection of ketamine-xylazine can effectively anesthetize different strains of mice for surgical procedures ^15, 35^. Also, mice receiving a subcutaneous injection of ketamine-xylazine had lower mortality than those receiving intraperitoneal injection ^15^.

In the present study, we compared the impacts of ketamine-xylazine, chloral hydrate, and isoflurane anesthetic agents on the surgical, behavioral, and histological parameters of the rat MCAO model. According to our findings, the CH group represented the highest, while Ket-Xyl/ SC showed the lowest mortality rate, up to three days after cerebral ischemia. The influence of anesthetic regimes on the mortality rate in MCAO models has been evaluated in several studies. In this regard, Bleilevens et al. (2013) have reported that isoflurane anesthesia reduced the first 24h mortality rate in MCAO rats compared to Ket-Xyl/ IP ^38^. While, Maud and colleagues found no significant difference in 24□h mortality rate between the isoflurane and chloral hydrate anesthesia groups ^13^. We also compared the effects of different injectable and inhalational anesthetics on the immobilization, induction, and recovery times. Isoflurane inhalation anesthesia led to faster induction and rapid recovery than injectable groups; however, there was no significant difference between injectable anesthesia groups.

Neurological deficits, infarction, and cognitive impairments are the main consequences of cerebral ischemia. In the present study, we found that 30 min MCAO induced neurological deficits, 1 and 3 days after the surgery. Moreover, our data confirmed the brain infarcts, three days after MCAO. Interestingly, our results showed no significant differences in the infarct size and neurological deficits among experimental groups receiving different anesthetic protocols. To date, evidence regarding the effects of anesthesia on infarct size and behavioral outcomes in animal models of stroke is controversial. Maud et al. have reported no significant difference in the infarct size between isoflurane and chloral hydrate anesthetized rats after 60 min MCAO ^13^. Similarly, Linou et al. have observed no significant difference in the infarct size between isoflurane and Ket-Xyl/ IP anesthetized mice, up to 7 days after MCAO ^39^. In contrast, Bleilevens et al. have discovered different infarct patterns in the isoflurane and Ket-Xyl/ IP anesthetized rats after 60 min MCAO. Striatal infarcts were detected more often in the isoflurane group, while cortical infarcts were found more frequently in the Ket-Xyl/ IP group at 24 h after reperfusion ^38^. Furthermore, Chen et al. have reported that the use of isoflurane or Ket-Xyl/ IP had similar effects on the infarct size in mice; however, Ket-Xyl/ IP significantly attenuated neurological deficit scores after stroke ^40^. Conflicting results on the stroke outcomes are likely related to animal species, duration of MCAO or reperfusion, the dose of the anesthetic drugs and duration of anesthesia.

In the present study, we also showed that 30 min MCAO, regardless of the anesthetic regimens, could not impair spatial working memory, three days after reperfusion. In humans, cognitive impairments usually occur in the acute phase of stroke and may persist till sub-acute and chronic phases ^41^. In animal models of stroke, it has been reported that cognitive deficits induced by MCAO are more pronounced in passive avoidance task, one week after 60 min MCAO ^42^. Moreover, as shown in Tables 2-4, MCAO-induced memory impairments in rats are usually found in prolonged MCAO and in extended time windows after reperfusion.

## Conclusion

In summary, our finding showed that 30 min MCAO, regardless of the anesthetic regimens, can remarkably induce cerebral infarction along with moderate neurological deficits. In this approach, spatial working memory is not affected, which might be valuable when spatial memory impartment is not desirable. In addition, we propose subcutaneous injection of ketamine/xylazine can be a promising regimen to induce anesthesia in the rat model of MCAO with several advantages such as low mortality, cost-effectiveness, safety, ease of administration, and not requiring specialized equipment.

## Acknowledgements

This study was financially supported by Shiraz University of Medical Sciences (Grant number: 23902). Partial support was also provided by Prof. Dr. Inga D Neumann from the University of Regensburg, Germany and FAPESP (Grant numbers: 2019/12526-3 and 2016/25502-7).

## Box 1: Neurological deficits were graded as a scale of 0-4.

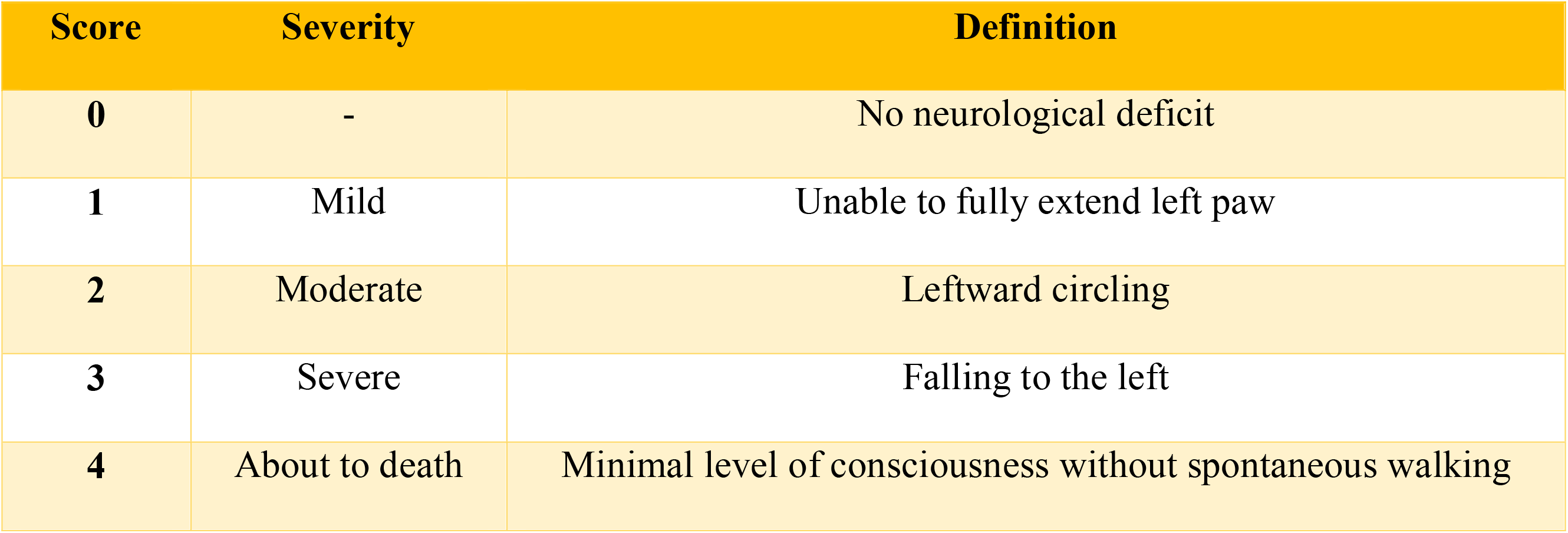

